# Differential gene expression analysis and physiological response characteristics of passion fruit (*Passiflora edulis*) buds under high-temperature stress

**DOI:** 10.1101/2022.09.07.506887

**Authors:** Hongli Wang, Jiucheng Zhao, Miao Lai, Yingqing Zhang, Wenwu Qiu, Yanyan Li, Hailian Tu, Qichang Ling, Xinfeng Fu

**Affiliations:** Qinzhou Branch of Guangxi Academy of Agricultural Sciences/Qinzhou City Institute of Agricultural Sciences,Qinzhou 535000,Guangxi,China; Institute of Horticulture, Guangxi Academy of Agricultural Sciences,Nanning 530007,Guangxi,China

**Keywords:** Passion fruit, High-temperature stress, Chlorophyll fluorescence, RNA-seq, differential expression gene, pathway

## Abstract

High temperature in summer is an unfavorable factor for passion fruit (Passiflora edulis), which can lead to restricted growth, short flowering period, few flower buds, low fruit setting rate, severe fruit drop, and more deformed fruit. To explore the molecular physiology mechanism of passion fruit responding to high-temperature stress, we use ‘Zhuangxiang Mibao’, a hybrid passion fruit cultivar, as the test material. Several physiological indicators were measured and compared between high-temperature (average temperature 38°C) and nonmoral temperature (average temperature 25°C) conditions, including photosynthesis, chlorophyll fluorescence parameters, POD, SOD activity and malondialdehyde content. We performed RNA-seq analysis combined with biochemistry experiment to investigate the gene and molecular pathways that respond to high-temperature stress. The results showed that some physiological indicators in the high-temperature group, including the net photosynthetic rate, stomatal conductance, intercellular CO_2_ concentration, transpiration rate, and the maximum chemical quantum yield of PS∥, were significantly lower than those of the control group. Malondialdehyde content was substantially higher than the control group, while superoxide dismutase and superoxide dismutase activities decreased to different degrees. Transcriptome sequencing analysis showed that 140 genes were up-regulated and 75 genes were down-regulated under high-temperature stress. GO and KEGG annotation analysis of differentially expressed genes revealed many metabolic pathways related to high-temperature stress. Further investigation revealed that 30 genes might be related to high-temperature stress, such as *CAO, GSH, WRKY*, and *HSP*, which have also been reported in other species. The results of real-time fluorescence quantitative PCR and RNA-seq of randomly selected ten genes are consistent, which suggests that the transcriptome sequencing results were reliable. Our study provides a theoretical basis for the mechanism of passion fruit response to high-temperature stress. Also, it gives a theoretical basis for the subsequent breeding of new heat-resistant passion fruit varieties.

## INTRODUCTION

Passion fruit is a perennial evergreen vine-like fruit tree. It was considered a tropical high-quality fruit tree with excellent development prospects based on its high nutritional, ornamental, and medicinal value (*Wang et al.,2022*). At the same time, passionflower is also a typical economic crop with the property of ‘easy to grow and difficult to manage’. One of the main reasons for the “difficulty to manage” is that heat damage caused by environmental changes often occurs and causes devastating damage to the industry. The optimal growth temperature of passionflower is 20°C-30°C. Above 30°C, the photosynthesis will decrease. If the temperature exceeds 35°C, a series of physiological lesions will appear in the plant, such as abnormal flower organs, leaves, fruit tissue, deformity, necrosis, and other sunburn symptoms. And the fruit set rate dropped to 10% to 15% of normal temperature. High temperatures not only change the phenotypic traits of the fruit but also cause the imbalance of cell homeostasis in the plant body and inhibit growth and development. High temperature also leads to large-scale dead seedlings, seriously affecting their quality and yield and causing enormous economic losses.

With the intensification of the greenhouse effect, the high-temperature stress caused by global warming has increasingly become a severe challenge for the modern agricultural production system. It is of great scientific significance and practical urgency to study the mechanism of heat regulation of plants. The research on plant heat resistance has become a hot research field in agricultural scientific research at home and abroad(*Chen, Guo et al.,2013;Akhoundnejad, Dasgan, Karabiyik,2020;Ma,2021;Ren,2021*). However, the research on the response of passion fruit to high-temperature heat damage began relatively late. Recent studies revealed the effect of high temperature on yield, quality, growth and development, and the effects of physiological and biochemical characteristics., however the molecular mechanism of passion fruit response to high-temperature stress is still unknown(*Tian et al.,2020;Tian et al.,2021;Huanget al.,2021;Lan, Shi, Li .2021*).

The emergence and development of RNA-seq technology have greatly accelerated research at the molecular level of plant science and provided new methods and ideas for a better understanding of the mechanism of heat resistance regulation in passion fruit. This technology is increasingly used to construct transcriptional maps under plant stress and screen plant stress-responsive genes. Therefore, it is possible to use RNA-seq technology to explore the heat-resistant response mechanism of passion fruit under high-temperature stress. This technology has revealed the high-temperature stress response mechanism of many plants. Huang Guiyuan et al. performed a transcriptome sequencing analysis of Vitis vinifera under high-temperature stress and screened BI1 with higher expression fold changes induced by high-temperature stress. By constructing a fusion expression vector to analyze the subcellular localization of *BI1* and using yeast two-hybrid system and other technical analyses, they found that *BI1* was a positive regulator in regulating Vitis vinifera response to high-temperature stress(*Huang et al.,2022*). A previous study by Chen Xueqian et al. shows that many *HSP* family genes are highly up-regulated when Cucumis sativus is subjected to high-temperature stress, and *AP2, MYB, WRKY, bHLH*, and *HSF* transcription factors show different expression patterns under high-temperature stress(*Chen, Han, Ren.2021*). In addition to conventional crops, the heat tolerance mechanisms of medicinal plants such as Pinellia ternata,Mentha canadensis,and Artemisia annua have also been revealed by transcriptome sequencing (*Lu et al.,2018;Da, Li, Gao. 2020;Guo et al.,2022*).

Therefore, to investigate the molecular regulation mechanism of passion fruit in response to high-temperature stress,we selected the high-temperature-sensitive passionflower variety ‘Zhuangxiang Mibao’ cultivated locally in Qinzhou, Guangxi,as the test material. By analyzing the physiological characteristic under high-temperature and normal temperature conditions, combined with transcriptome sequencing technology,the essential functional genes and regulatory factors involved in the high-temperature response of passion fruit were identified. Our study provides essential theoretical and practical significance to improve the tolerance to high temperatures and cultivate new varieties of passion fruit with high-temperature resistance through a genetic approach.

## MATERIAL AND METHODS

### Plant material

The test material ‘Zhuangxiang Mibao’ was planted in the Jiulong Experimental Base of Agricultural Science Research Institute, Qinzhou, Guangxi, in February 2022. Two hundred grafted seedlings were grown in the open air or experimental greenhouse, where the average daily temperature in May was 25° C and 38 °C, respectively. The young flower buds of ‘Zhuangxiang Mibao’ with the same experimental field, growth potential, external morphological characteristics, and seedling height of about 150 cm were selected. The surface of selected young flower buds was rinsed with 75% ethanol, dried with absorbent paper, and aliquoted with 100 mg per capped storage tube. The capped storage tube was quickly frozen in liquid nitrogen and stored at -80°C. Three biological replicates were set and numbered for six samples to filter false positive signals.

### Determination of Physiological index

To ensure the accuracy of photosynthetic index determination, the leaves with the same position, moisture, nutritional status, without insect spots, and disease spots were selected, and the experiments were performed within 9:00-11:30 in the morning when the photosynthesis induction period reached a steady state. A portable photosynthesis instrument (LI-6400XT) was used in the experimental base to determine the net photosynthetic rate (Pn), stomatal conductance (Gs), concentration of intercellular CO_2_ (Ci) and transpiration rate (Tr) and other photosynthetic indicators.After changing the fluorescent leaf chamber probe, the chlorophyll fluorescence parameter values were measured, including dark-adapted maximum fluorescence yield (Fm), dark-adapted minimum fluorescence yield (Fo), light-responsive maximum fluorescence yield (Fm’), the minimum fluorescence yield of photoreaction (Fo’) and the steady-state fluorescence yield (Fs). The photochemical potential activity of PSII (Fv/Fo) and the maximum photochemical quantum efficiency of PSII (Fv/Fm) were calculated based on the chlorophyll fluorescence parameters.Peroxidase activity(POD),superoxide dismutase (SOD) activity, and malondialdehyde (MDA) content were detected indoors using biochemical kits (Solarbio, Beijing). Measured data were analyzed using DPS v1.9.2 and Excel 2022.

### Total RNA extraction, quality inspection, cDNA library construction and sequencing

3 μg total RNA was extracted from each flower bud sample using Tiangen Polysaccharide and Polyphenol Kit (QIAGEN, Germany). After passing the quality inspection, Illumina Novaseq 6000 (sequencing read length is PE150) was used for library construction and RNA-Seq sequencing.

### Analysis of sequencing data

To ensure the quality and reliability of data analysis, high-quality clean data (clean reads) were obtained after removing errors, or low-quality reads from the raw sequence reads (Raw reads) obtained by sequencing. The reference genome sequence and annotation file of passion fruit were downloaded from the National Genome Data Center (NGDC) of China, project number PRJCA004251 (https://ngdc.cncb.ac.cn/gwh/Assembly/17982/show) (*Xia et al.,2021*). The clean reads were compared with the reference genome of passion fruit using HISAT2 software to obtain the information of reads on the reference genome(*Mortazavi et al.,2008*).

### Gene expression levels and differentially expressed gene enrichment analysis

Considering the influence of sequencing depth and gene length on the count of fragments, the FPKM value was used to estimate the gene expression level. Based on the negative binomial distribution, the DESeq2 software was used to screen differentially expressed genes (DEGs) with the criteria of Padj ≤0.05 (Padj is the corrected P) and |log2(Fold Change)|≥1. GO function and KEGG pathway enrichment analysis were performed on the differential gene sets using the cluster Profiler(*Yu et al.,2012*).

### Validation of Target gene using qRT-PCR

The complementary DNA (cDNA) was prepared by reverse transcription from cryopreserved passion fruit bud RNA using the cDNA Synthesis Kit (Thermo). The expression of EF1 and Ts in each tissue is relatively stable, so EF1 and Ts were used as positive controls(*Zhao et al.,2022;Tu,Fan,Xu.2022*).OLIGO was used to design specific primers for differentially expressed genes of passion fruit related to high-temperature stress.The primer sequences were delivered to Sangon Bioengineering (Shanghai) for synthesis (Table 1). The qRT-PCR assay was performed in a 20 μL of system: 0.4 μl of upstream and downstream primers (10 μmol·L−1), 1.0 μl of cDNA template, 10.0 μl of 2×SG Green qPCR Mix, 0.4 μl of ROX, and 7.8 μl of ddH_2_O. Amplification procedure: pre-denaturation at 95°C for 10 min; denaturation at 95°C for 15s, annealing at 55°C for 15s, renaturation at 60°C for 30s, extension at 72°C for 45s, a total of 45 cycles. The melting curve acquisition program was: heating to 95 °C for 15 s, dropping to 60 °C for 30 s, and heating to 95 °C for 15 s. The amplification process was conducted in StepOne™ Plus Real-Time PCR System (Applied Biosystems). The expression levels of differential genes were calculated by the 2^-△△CT^ method(*Tatarowska et al.,2020*).

**Table 1.**
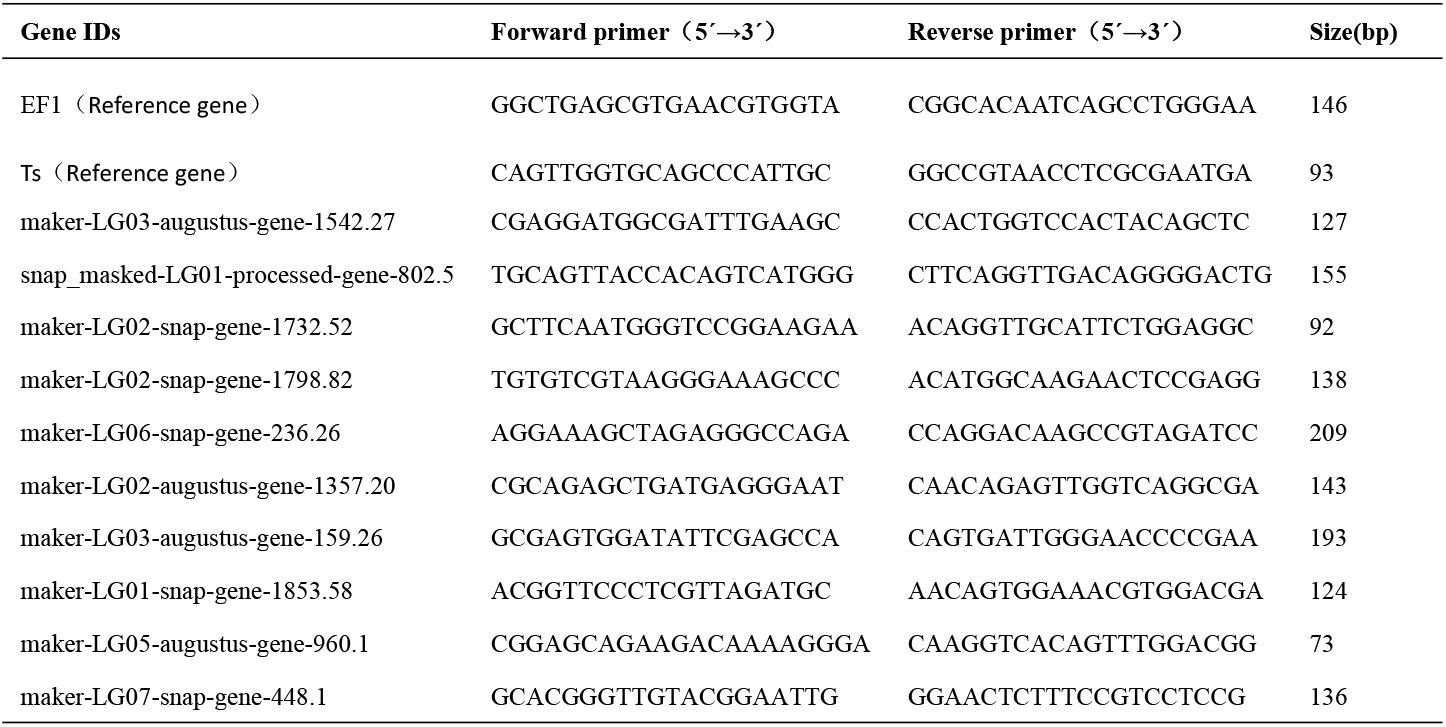
Real-time quantitative fluorescence gene primer sequences.

### Data availability

The transcriptome sequence and annotation information of passion fruit buds measured in this study have been uploaded to the National Gene Bank of China (CNGB) and the National Center for Biological Information (NCBI) in the United States for relevant researchers to consult.

## RESULTS

### Changes of physiological and biochemical indexes of passion fruit in high-temperature conditions

The passion fruit undergoes a series of physiological and biochemical changes under high-temperature stress. The net photosynthetic rate, stomatal conductance, intercellular CO_2_ concentration, and transpiration rate of the high-temperature group were significantly lower than those of the control group (Table 2). Under high-temperature conditions, the passion fruit leaves quickly lose water. When the water deficit is severe, the content of abscisic acid in the leaves increases rapidly, which causes the stomata to close, and the stomatal conductance and photosynthetic rate decrease accordingly. In the healthy physiological state, the maximum photochemical quantum efficiency value of PSII of the passion fruit is 0.815. However, the lowest measured value under high-temperature stress is 0.545, indicating that photosynthesis is affected and the poor health status. POD activity, SOD activity, and MDA content are critical indicators to measure the plant stress response. Among them, the MDA content is one of the essential indicators for judging the degree of damage to the plant cell membrane. Long-term high-temperature stress led to a significant increase in the MDA content in the leaves of passion fruits, and the POD and SOD activities showed varying degrees of change.

**Table 2.** Changes of main physiological characteristics of passionflower under high-temperature stress.

| Category | Pn($\mu\text{mol CO}_2\text{m}^{-2}\text{s}^{-1}$) | Gs($\text{molH}_2\text{O m}^{-2}\text{s}^{-1}$) | Ci($\mu\text{mol CO}_2\text{ mol}^{-1}$) | Tr($\text{mmolH}_2\text{Om}^{-2}\text{s}^{-1}$) | F<sub>v</sub>/F<sub>o</sub> | F<sub>v</sub>/F<sub>m</sub> | POD (U/$\mu\text{g}$) | SOD (U/$\mu\text{g}$) | MDA (nmol/g) |
| --- | --- | --- | --- | --- | --- | --- | --- | --- | --- |
| HT<sub>1</sub> | 5.041 | 0.051 | 218.262 | 1.269 | 2.539 | 0.584 | 998.628 | 37.994 | 19.358 |
| HT<sub>2</sub> | 2.139 | 0.025 | 243.179 | 0.657 | 2.755 | 0.612 | 1569.261 | 17.532 | 13.012 |
| HT<sub>3</sub> | 4.675 | 0.048 | 219.228 | 1.203 | 2.699 | 0.545 | 3851.824 | 36.435 | 12.373 |
| CK<sub>1</sub> | 14.929 | 0.330 | 307.844 | 7.016 | 3.781 | 0.797 | 285.323 | 23.908 | 10.655 |
| CK<sub>2</sub> | 19.431 | 0.295 | 271.063 | 6.598 | 4.242 | 0.815 | 570.641 | 35.929 | 8.923 |
| CK<sub>3</sub> | 16.424 | 0.305 | 293.871 | 6.742 | 3.643 | 0.772 | 2853.26 | 17.161 | 6.881 |

### Transcriptome sequencing analysis

The flower bud samples of the high-temperature group (HT_1_, HT_2_, and HT_3_) and the control group (CK_1_, CK_2_, and CK_3_) were sequenced, and a total of 41.31 G Clean Data was obtained, and each sample produced an average of more than 6.89 Gb of high-quality data. The percentage of Q20 bases and Q30 bases in each sample was greater than 97.85% and 93.84%, respectively, and the GC content ranged from 44.92% to 45.55%. The mapping rates of sequenced genes ranged from 77.71% to 86.10%, and an average of 74.49% of the reads could be aligned to the unique position in the genome (Table 3). The overall quality of the six samples met the requirements for subsequent analysis.

**Table 3.**
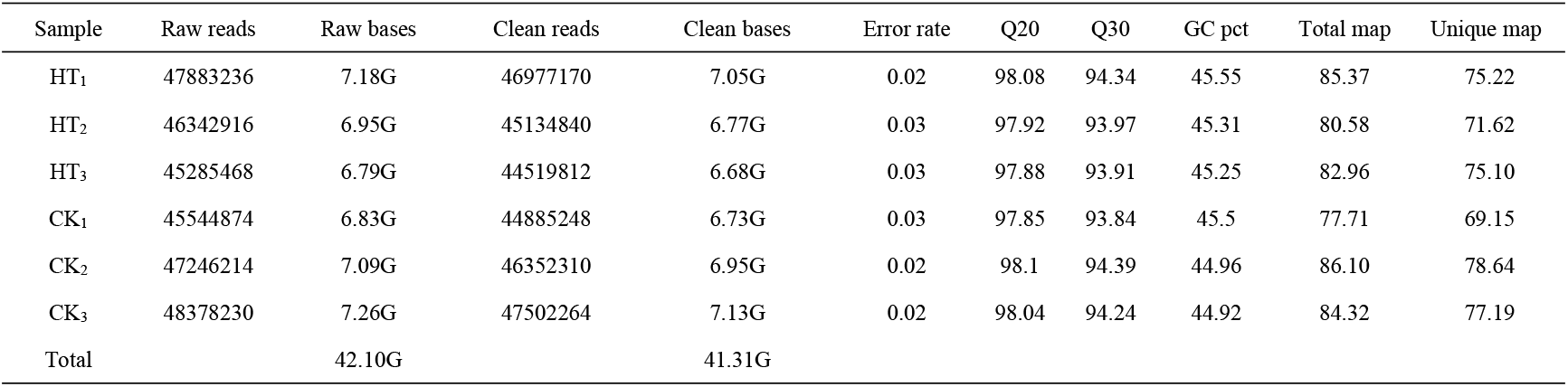
Statistical table of sample sequencing data evaluation.

| Sample | Raw reads | Raw bases | Clean reads | Clean bases | Error rate | Q20 | Q30 | GC pct | Total map | Unique map |
| --- | --- | --- | --- | --- | --- | --- | --- | --- | --- | --- |
| HT<sub>1</sub> | 47883236 | 7.18G | 46977170 | 7.05G | 0.02 | 98.08 | 94.34 | 45.55 | 85.37 | 75.22 |
| HT<sub>2</sub> | 46342916 | 6.95G | 45134840 | 6.77G | 0.03 | 97.92 | 93.97 | 45.31 | 80.58 | 71.62 |
| HT<sub>3</sub> | 45285468 | 6.79G | 44519812 | 6.68G | 0.03 | 97.88 | 93.91 | 45.25 | 82.96 | 75.10 |
| CK<sub>1</sub> | 45544874 | 6.83G | 44885248 | 6.73G | 0.03 | 97.85 | 93.84 | 45.5 | 77.71 | 69.15 |
| CK<sub>2</sub> | 47246214 | 7.09G | 46352310 | 6.95G | 0.02 | 98.1 | 94.39 | 44.96 | 86.10 | 78.64 |
| CK<sub>3</sub> | 48378230 | 7.26G | 47502264 | 7.13G | 0.02 | 98.04 | 94.24 | 44.92 | 84.32 | 77.19 |
| Total |  | 42.10G |  | 41.31G |  |  |  |  |  |  |

### Screening of differentially expressed genes

We performed differential expression analysis based on the sequencing results to find the gene response to high-temperature stress in passion fruit. Hierarchical clustering was used to perform bidirectional cluster analysis based on the FPKM values of the genes (Fig. 2A). Statistical analysis was performed on the expression of the differentially expressed genes between the high-temperature group and the control group, and a total of 215 significantly differentially expressed genes were screened. Among them, 140 genes were up-regulated and 75 genes were down-regulated. The up-regulated genes were about twice as many as the down-regulated genes (Fig. 2B). Multiple genes showed differential expression patterns when passion fruit responded to high-temperature stress, indicating that the response to high-temperature stress was a complex process involved in multi-gene interaction.

**Figure 1.**
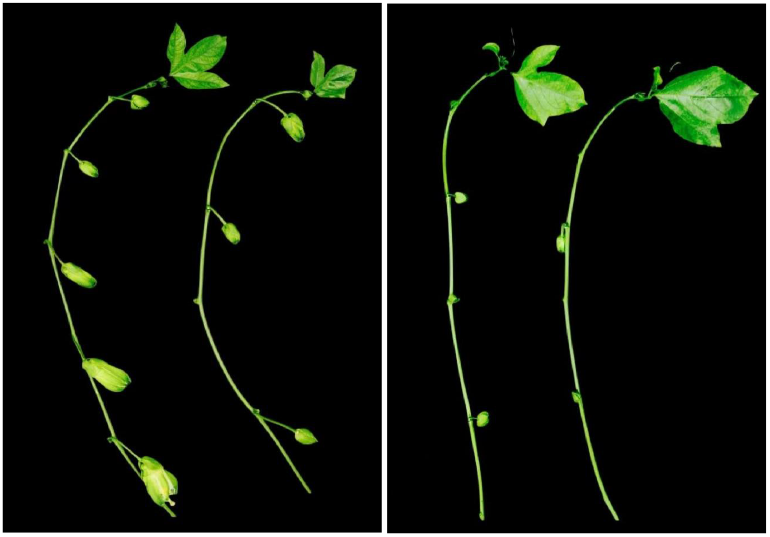
Comparison of flower bud number between secondary and tertiary tendrils. A. Number of secondary vine buds B. Number of tertiary vine buds (Left CK group, right HT group)

**Figure 2.**
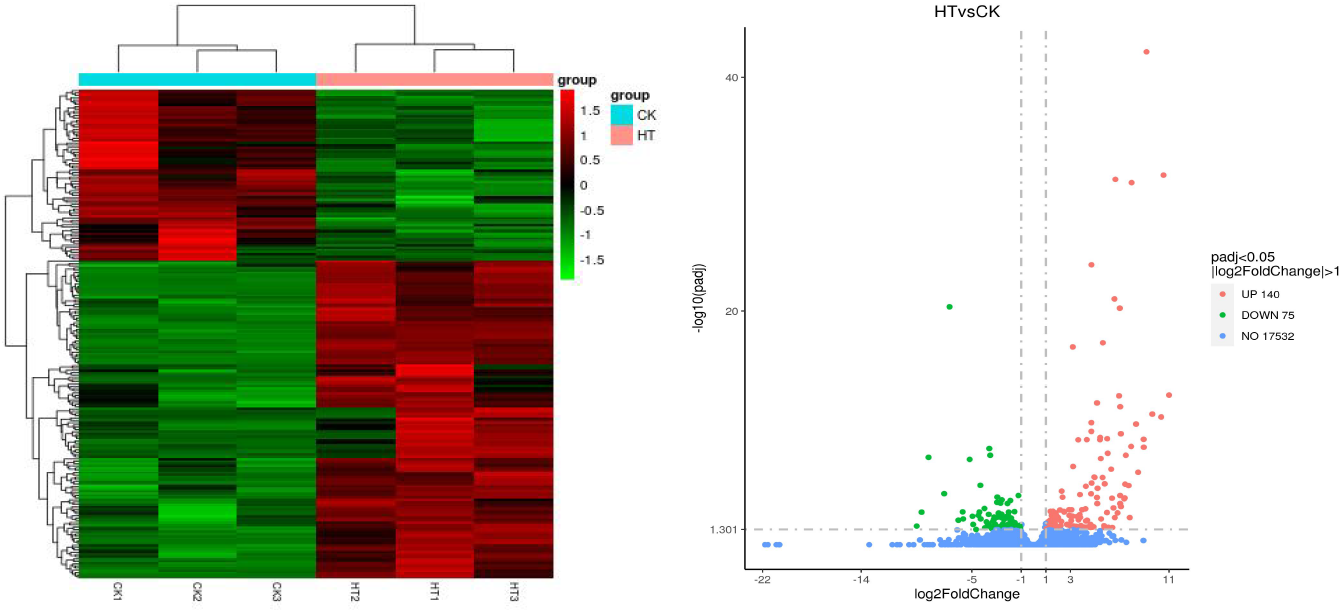
Cluster map, heat map and volcano map of differentially expressed genes.

### GO enrichment analysis of DEGs

GO enrichment analysis was used mainly to predict the function of differentially expressed genes. To further analyze the functions of differentially expressed genes, we performed GO function annotation analysis of 215 DEGs. We found that 195 DEGs were annotated into 331 categories, of which 134 were categories belonging to the biological process, 71 categories belonging to cellular components, and 126 categories belonging to molecular functions. Among them, 131 DEGs were up-regulated, and 64 DEGs were down-regulated. Figure 3 shows the GO terms in which DEGs are significantly enriched in the GO data. In terms of biological processes, in addition to carbohydrate metabolism (9, 13.04%, GO: 0005975) was the most significant enrichment, there were also stimulation responses (5, 7.24%, GO: 0050896), transmembrane transport (4, 5.80%, GO: 0055085), regulating cellular metabolic processes (4, 5.80%, GO: 0031323) and stress response (3, 4.35%, GO: 0006950) and other related stress-related factors GO term, indicating that the high-temperature has affected the normal life activities of passion fruit. In terms of cellular components, DEGs were mainly enriched in protein complexes (8, 33.33%, GO: 0032991), membrane-bound organelles (6, 25%, GO: 0043227), intracellular organelles (5, 20.83%, GO: 0044446) and other GO terms. In terms of molecular functions, DEGs are mainly involved in hydrolase activity (9, 8.82%, GO: 0004553), enzymatic activity regulation (6, 5.88%, GO: 0030234) and oxidoreductase activity (6, 5.88 %, GO: 0016705). These results suggest that high-temperature stress has impaired the metabolic reaction rate and normal physiological activities in cells. The cell can adapt to the changes in external conditions by regulating enzyme activity and maintaining its life activity.

**Figure 3.**
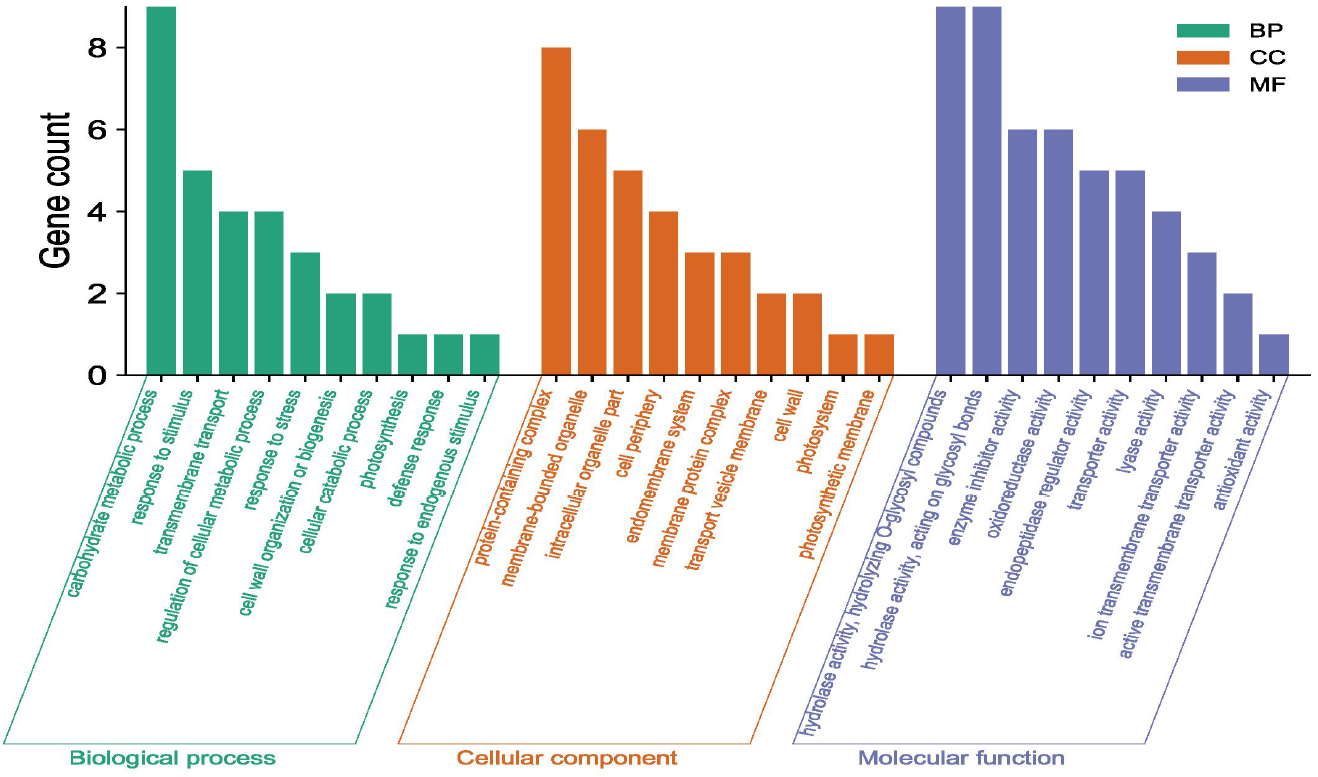
GO functional classification analysis of differentially expressed genes KEGG classification and pathway enrichment analysis of DEGs.

To clarify the biological metabolic pathways of passion fruit in response to high-temperature stress, we performed a metabolic pathway regulatory network enrichment analysis of DEGs between the high-temperature group and the control group. A total of 53 DEGs were annotated to 52 KEGG metabolic pathways, including metabolism (34), genetic information processing (10), environmental information processing (4), cellular process (2), and biological system (2). Among them, 34 DEGs were up-regulated and 19 DEGs were down-regulated. Figure 4 shows the important metabolic pathways with the highest degree of DEGs enrichment. The five metabolic pathways with a large number of DEGs annotations are starch and sucrose metabolism (5, 9.43%, ID: 00500), ribosome (5, 9.43%, ID: 03010), fatty acid metabolism (4, 7.55%, ID: 01212), amino sugar and nucleotide sugar metabolism (4, 7.55%, ID: 00520), and plant hormone signal transduction (3, 5.66%, ID: 04075). Based on the P value, we take a further metabolic pathway enrichment analysis of the up-regulated and down-regulated DEGs. It was found that the up-regulated DEGs in passion fruit buds were mainly related to monoterpenoid biosynthesis, amino sugar and nucleotide sugar metabolism, pyruvate metabolism, plant hormone signal transduction, porphyrin and chlorophyll metabolism, and photosynthesis under high-temperature condition (Fig. 5A). DEGs with down-regulated expression are mainly enriched in sesquiterpene and triterpenoid biosynthesis, starch and sucrose metabolism, glutathione metabolic pathways, amino acid metabolism, unsaturated fatty acid biosynthesis, and MAPK signaling (Fig. 5B). Among them, plant hormone signal transduction, starch and sucrose metabolism pathways, porphyrin and chlorophyll metabolism, and photosynthesis pathways all play essential biological functions in response to a variety of plant stress, which suggested that these metabolic pathways may be involved in response to high-temperature stress.

**Figure 4.**
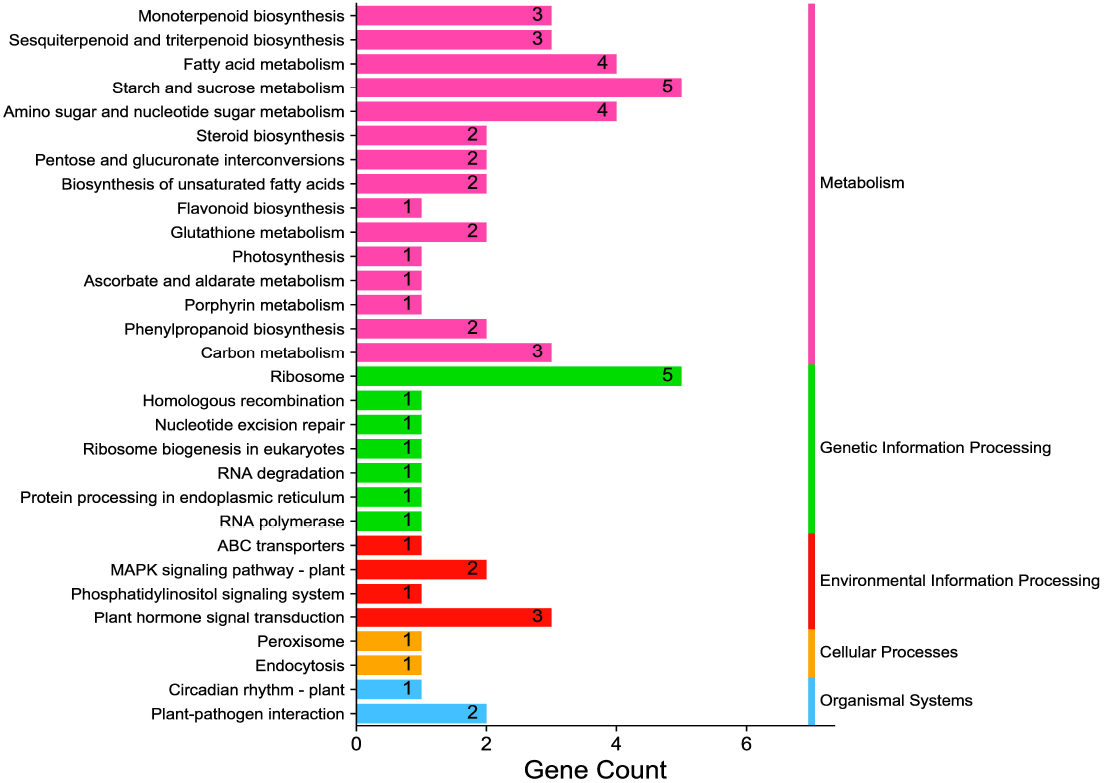
KEGG enrichment pathway of differentially expressed genes.

**Figure 5.**
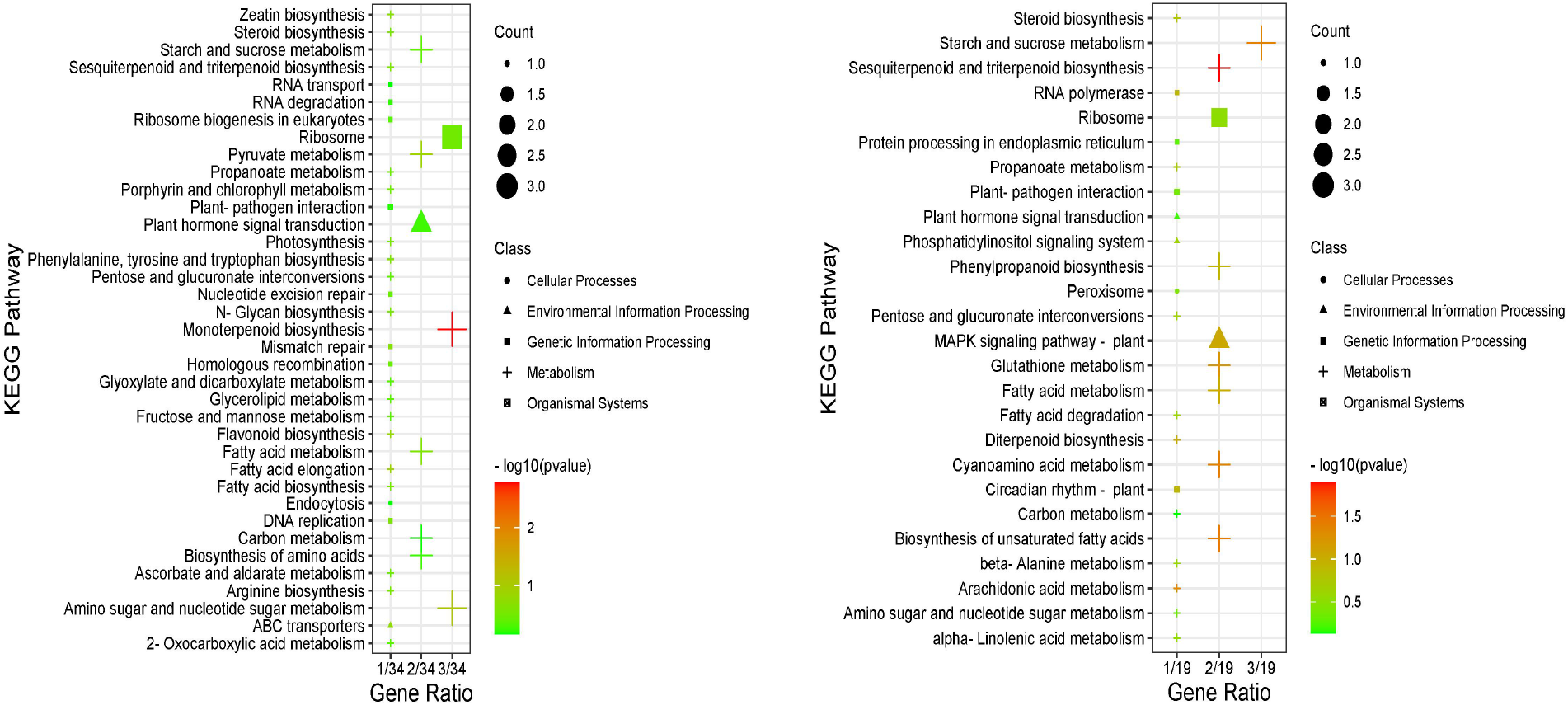
KEGG pathway enrichment analysis. A:Results of up-regulating DEGs pathway enrichment B: Enrichment results of DEGs pathway down-regulated by expression level

### Candidate genes related to high temperature stress

Through the comparison of differentially expressed genes between the high-temperature group and control group, followed by GO classification and KEGG enrichment analysis, and combined with relevant literature, we screened out 28 candidate genes that may be involved in high-temperature stress in passion fruit. These genes involved in photosynthesis, phytohormone, glutathione, protein kinase, heat shock proteins, nighttime lighting and clock regulation, and thioredoxin (Table 4)(*Wang, 2021;Hao,2021;Chen et al.,2020*).

**Table 4.** Differential expression of 28 candidate genes in passionflower under high-temperature stress.

| Gene IDs | Gene name | Gene expression (FPKM) |  | log<sub>2</sub> (Fold Change) |
| --- | --- | --- | --- | --- |
|  |  | HT | CK |  |
| Photosynthesis |  |  |  |  |
| maker-LG06-snap-gene-236.26 | PesCAO1 | 41.52 | 14.22 | 1.55 |
| augustus_masked-LG06-processed-gene-154.56 | PesCAO2 | 28.86 | 0.10 | 8.33 |
| maker-LG03-snap-gene-1549.4 | PesCAO3 | 16.38 | 0.63 | 4.70 |
| maker-LG01-snap-gene-1050.8 | PesCAO4 | 10.76 | 0.43 | 4.70 |
| maker-LG02-augustus-gene-1357.20 | PesCAO5 | 2.60 | 293.76 | -6.82 |
| snap_masked-LG06-processed-gene-149.9 | PesCAO6 | 4.42 | 0.04 | 6.95 |
| Plant hormone signal |  |  |  |  |
| snap_masked-LG01-processed-gene-802.5 | PesSAUR1 | 28.71 | 1.90 | 3.95 |
| maker-LG02-snap-gene-1732.52 | PesXTH1 | 15.06 | 1.66 | 3.19 |
| maker-LG02-snap-gene-1798.82 | PesPP1 | 3.03 | 17.07 | -2.49 |
| Glutathione |  |  |  |  |
| maker-LG03-augustus-gene-1542.27 | PesGST1 | 0.62 | 13.68 | -4.47 |
| maker-LG07-snap-gene-228.31 | PesGST2 | 11.81 | 5.03 | 1.23 |
| maker-LG06-snap-gene-57.0 | PesGSH1 | 24.31 | 105.87 | -2.12 |
| Protein kinase |  |  |  |  |
| maker-LG03-augustus-gene-159.26 | PesPK1 | 40.64 | 7.35 | 2.47 |
| maker-Contig20-augustus-gene-1.8 | PesPK2 | 23.27 | 0 | 8.96 |
| maker-Contig29-augustus-gene-1.94 | PesPK3 | 7.75 | 3.03 | 1.35 |
| maker-LG03-augustus-gene-151.65 | PesPK4 | 4.31 | 0.54 | 3.04 |
| maker-Contig5-snap-gene-4.19 | PesPK5 | 1.84 | 0.14 | 3.75 |
| augustus_masked-LG01-processed-gene-1682.46 | PesPK6 | 1.47 | 0.18 | 3.06 |
| maker-LG05-snap-gene-37.79 | PesPK7 | 0.86 | 0.04 | 4.63 |
| maker-LG02-snap-gene-1846.50 | PesPK8 | 0.11 | 1.68 | -3.94 |
| Other |  |  |  |  |
| maker-LG06-augustus-gene-1233.26 | PesHSP1 | 0.96 | 0.06 | 4.14 |
| maker-LG01-snap-gene-1801.51 | PesHSP2 | 6.72 | 133.56 | -4.31 |
| maker-LG07-snap-gene-448.1 | PesWRKY1 | 0.05 | 1.32 | -4.66 |
| maker-LG07-augustus-gene-269.0 | PesWRKY2 | 0.34 | 0 | 6.08 |
| maker-LG05-augustus-gene-960.1 | PesLNK1 | 175.87 | 47.40 | 1.89 |
| maker-LG01-augustus-gene-177.30 | PesLNK2 | 96.70 | 41.56 | 1.22 |
| maker-LG05-augustus-gene-227.12 | PesLNK3 | 5.80 | 1.61 | 1.86 |
| maker-LG01-snap-gene-1853.58 | PesTRX1 | 105.92 | 35.19 | 1.59 |

### Analysis of photosynthesis-related DEGs

Chlorophyll plays a role in photosynthesis and is essential for plant development processes and responses to environmental stimuli(*Li et al.,2021*). High-temperature stress leads to significant changes in photosynthesis-related energy metabolism and physiological processes. Compared to the control group, *PesCAO1, PesCAO2, PesCAO3*, and *PesCAO6* were up-regulated after high-temperature stress. It has been reported that *CAOs* were involved in generating reactive oxygen species, which are signals and responses to abiotic stress. In this study, the expression level of chlorophyll a oxygenase *PesCAO1* was 41.52 in the high-temperature group, which was 2.92 times that of the control group (14.22). These results suggested that the passion fruit is more likely to maintain a high photosynthetic capacity under high-temperature stress.

### Analysis of plant hormone-related DEGs

High-temperature stress can cause changes in hormone metabolism and signal transduction in plant cells. Transcriptome sequencing analysis showed that the expression of auxin-responsive protein *PesSAUR1* and xyloglucan endotransglucosylase *PesXTH1* were up-regulated, while the expression of protein phosphatase *PesPP1* was down-regulated under high-temperature stress. These results indicate that different plant hormone response genes exhibited different expression patterns under high-temperature stress.

### Analysis of glutathione-related DEGs

Plant cells will produce excess reactive oxygen species under abiotic stress, resulting in oxidative stress. As a ubiquitous antioxidant in plants, glutathione can eliminate excess reactive oxygen species and keep cells in the homeostasis of redox reactions. The glutathione level is closely related to the tolerance of plants in the face of adversity. In this study, three differentially expressed genes were enriched in the glutathione metabolic pathway, including the glutathione peroxidase (GSH) gene and the glutathione S-transferases (GST) gene. These results suggested that passion fruit reduces the peroxidation of lipids in cells by regulating the expression of genes related to the glutathione metabolic pathway, thereby improving the tolerance to high-temperature stress.

### Analysis of protein kinase-related DEGs

Protein kinases mainly catalyze the phosphorylation of proteins, which is involved in the transmission of intracellular signals of plants under high-temperature stress. We found eight protein kinase-related genes were up-regulated under high-temperature conditions. They may be involved in regulating phytohormone content, including auxin, abscisic acid, gibberellin, and cytokinin.

### Real-time fluorescence quantitative analysis

To validate the reliability of the transcriptome sequencing data, we randomly select 10 DEGs (six up-regulated genes and four down-regulated genes) and verified by qRT-PCR, including chloroplast genes (*PesCAO1, PesCAO5*), auxin-responsive protein gene *PesSAUR1*, Xyloglucan endotransglycosylase gene *PesXTH1*, protein phosphatase gene*PesPP1*, glutathione S-transferase gene *PesGST1*, protein kinase gene *PesPK1*, WRKY transcription factor gene *PesWRKY1*, night lighting and clock regulation gene *PesLNK1* and thioredoxin gene *PesTRX1* (Figure 6). Correlation analysis shows that the changing trend and fold between real-time quantitative PCR and transcriptome data are generally consistent, and the R^2^ is 0.9348, which confirms the reliability of the transcriptome sequencing results.

**Figure 6.**
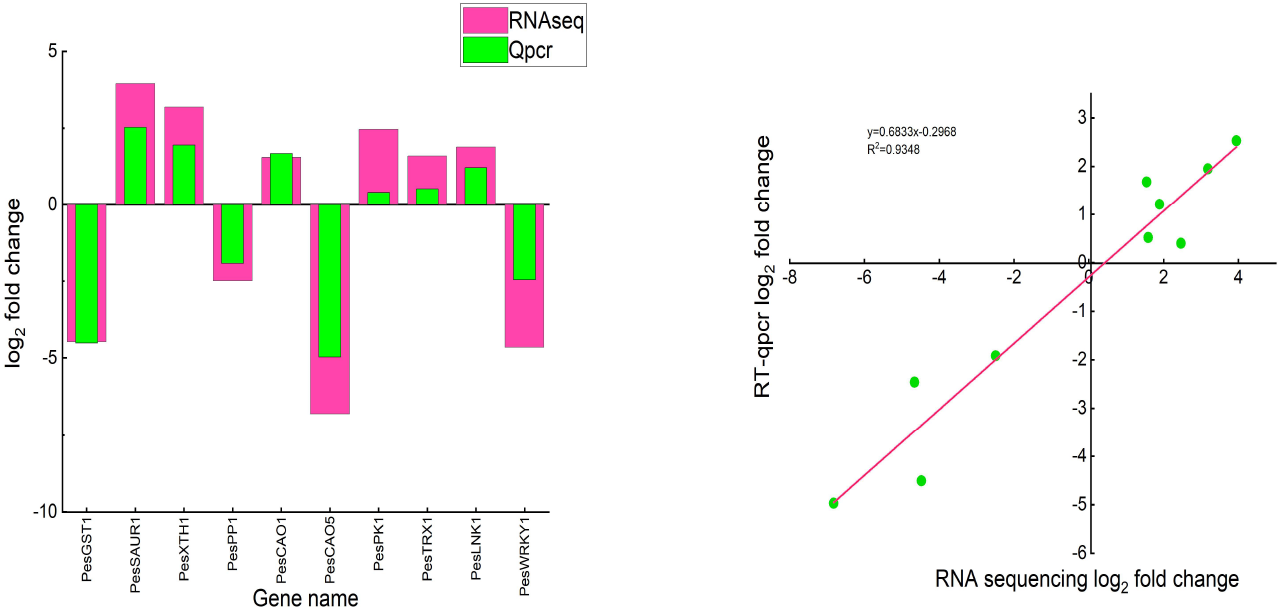
Real-time fluorescence quantitative results for differential genes.

## DISCUSSION

High temperature is one of the abiotic stress factors affecting crop production, which restricts the growth and development of crops. Studies have shown that high-temperature stress also changes the physiological and biochemical states, including decreased photosynthetic rate, blocked chlorophyll synthesis, and accumulated reactive oxygen species, which in turn causes a series of metabolic changes in cells. Once the plant cell membrane senses the high-temperature stress, cells initiate the transmission of signal molecules, causing changes in a series of life activities such as enzyme activity, cell structure, gene expression, and regulation to resist and ultimately alleviate the harm of high temperature(*Foyer, Noctor,2005;Han et al.,2010*). When the damage caused by the high-temperature stress exceeds the regulation ability of the organism, the plant begins to show the symptoms of heat damage in the form of appearance, such as the leaves will turn yellow or curling, the fruit will shrink and losing water, and the branches will wither and die.

Malondialdehyde content is an important indicator to evaluate the damage degree of the plant cell membrane system. In this study, the content of malondialdehyde in passion fruit was higher under high-temperature stress, indicating that the cell membrane system was damaged under high-temperature stress of 38 °C. A previous study shows that the MDA content in kiwifruit under high-temperature stress significantly increased, and the results were consistent with ours (*Dong,2018*).In this study,215 differentially expressed genes related to regulating high-temperature stress were obtained at the transcriptional level based on RNA-seq technology. GO functional annotation analysis showed that the differentially expressed genes under high-temperature stress were mainly enriched in the biological process database of carbohydrate metabolism, stimulus-response, transmembrane transport, regulation of cellular metabolism, and stress response metabolism. In the GO cellular component database, differentially expressed genes are mainly enriched in protein complexes, membrane-bound organelles, and intracellular organelles. In the GO molecular function database, it is primarily enriched in hydrolase activity, enzyme activity regulation, and oxidoreductase activity. These results show that the response to high-temperature stress is a complex physiological process, and the resistance to damage caused by high-temperature stress requires the cooperation of multiple biological processes, cellular components, and molecular functions. In the KEGG enrichment analysis, 34 pathways are enriched, indicating that passion fruit enhances various metabolic activities to resist the damage caused by high temperature. Another study reported that plant hormone signal transduction, glutathione metabolism, porphyrin and chlorophyll metabolism, MAPK signaling pathway, and photosynthesis pathway were involved in plant stress signal transduction(*Jiang et al.,2021; Zhou et al.,2022*). Our study also detected the high expression of differential genes in the metabolic pathways mentioned above, indicating that all of these signaling pathways respond to high-temperature stress in passion fruit. Our study also found that the heat shock proteins PesHSP1 and PesHSP2 are involved in response to high-temperature stress, which is consistent with the study using Lentinus edodes and strawberries (*Xin,2016;Zhang et al.,2022*). Based on transcriptome sequencing, many studies have been conducted on the differential expression of genes under heat stress in crops. Although many differentially expressed genes related to high-temperature stress have been identified, most research sites are limited to crop leaves. Matsuda H et al. pollinated passion fruit at different temperatures of 28-42 °C and then recorded the fruit set rate and seed number to determine the critical high temperature that negatively affected the fruit set rate (*Matsuda, Higuchi 2020a;Matsuda, Higuchi 2020b*). The study found that the fruit set rate decreased significantly when the temperature was greater than 38 °C. They also found that the pollen tube of all pollinated flowers, which were incubated at 28-32°C in vitro, was observed to reach the embryo sac within 24 hours. When the culture temperature was higher than 34°C, the shape of the pistils of all isolated flowers was disordered, and the pollen tube could not reach the embryo sac within 24h. When pollination at 40°C, the pollen still germinated on the stigma but did not extend to the style within 24 h. Passion fruit buds often encounter high-temperature during the bud differentiation period. High-temperature stress leads to a decrease in pollen vigor, which affects pollination and fruiting, resulting in a significant reduction in yield. Therefore, it is more targeted and innovative to choose the flower bud as the sequencing site to investigate the effect of high-temperature stress on the yield and quality of passion fruit.

## CONCLUSION

In this study, ‘Zhuangxiang Mibao’ was used as the test material. By measuring and analyzing the physiological indicators related to the photosynthetic system of leaves under different temperature conditions and combined with the transcriptome sequencing of the flower buds, we obtained some differentially expressed genes related to high-temperature stress. The functions of the differentially expressed genes were annotated. Combined with the physiological changes of leaves under high-temperature stress, candidate core gene groups, including CAO, GSH, WRKY, and HSP were screened out, which may be involved in the regulation of the process of high-temperature stress in passion fruit. The qRT-PCR analysis results of the ten differentially expressed genes were consistent with the results of RNA-seq, indicating that the transcriptome data were reliable. The changes in gene expression and physiological indicators were the same. Therefore, it is speculated that there may be correlations between the changes in physiological indexes and the expression of related genes under high-temperature stress. This study is a preliminary exploration of the response of passion fruit to high-temperature stress. Our transcriptome data will be used as a reference data for investigating passion fruit in response to high-temperature stress, such as molecular marker development, functional gene prediction, metabolic pathway exploration, and critical gene mining of high-temperature stress.

## Author contribution

Qichang Ling and Xinfeng Fu designed the project and guided the experimental design, data analysis, and thesis revision. Hongli Wang designed and performed most of the experiment, completed the data analysis, and wrote the manuscript. Jiucheng Zhao,Miao Lai,Yingqing Zhang, Wenwu Qiu,Yanyan Li and Hailian Tu participated in experimental design, performed experiments, assisted in data analysis, and revised the thesis. All authors read and agree to the final manuscript.

---

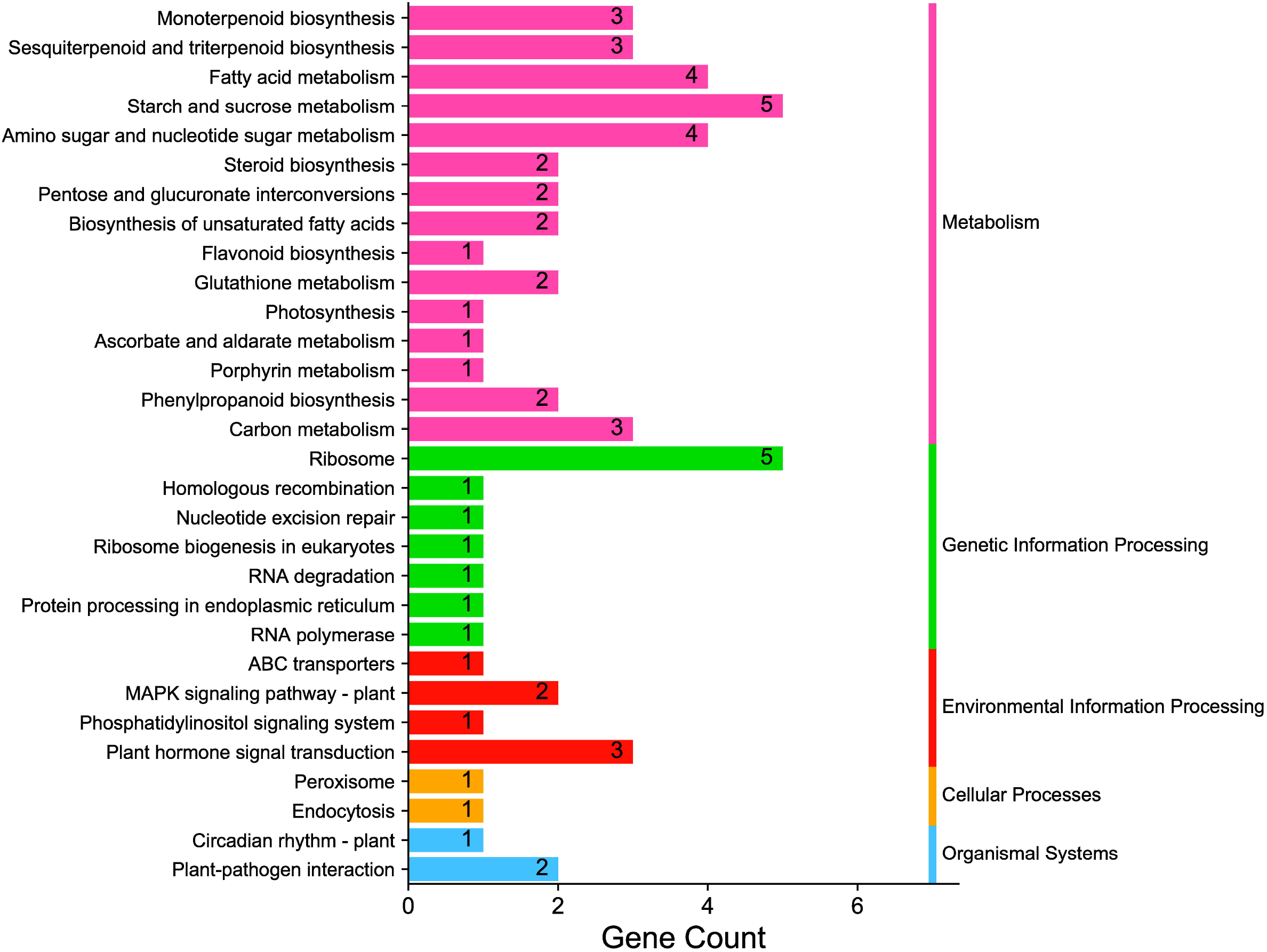

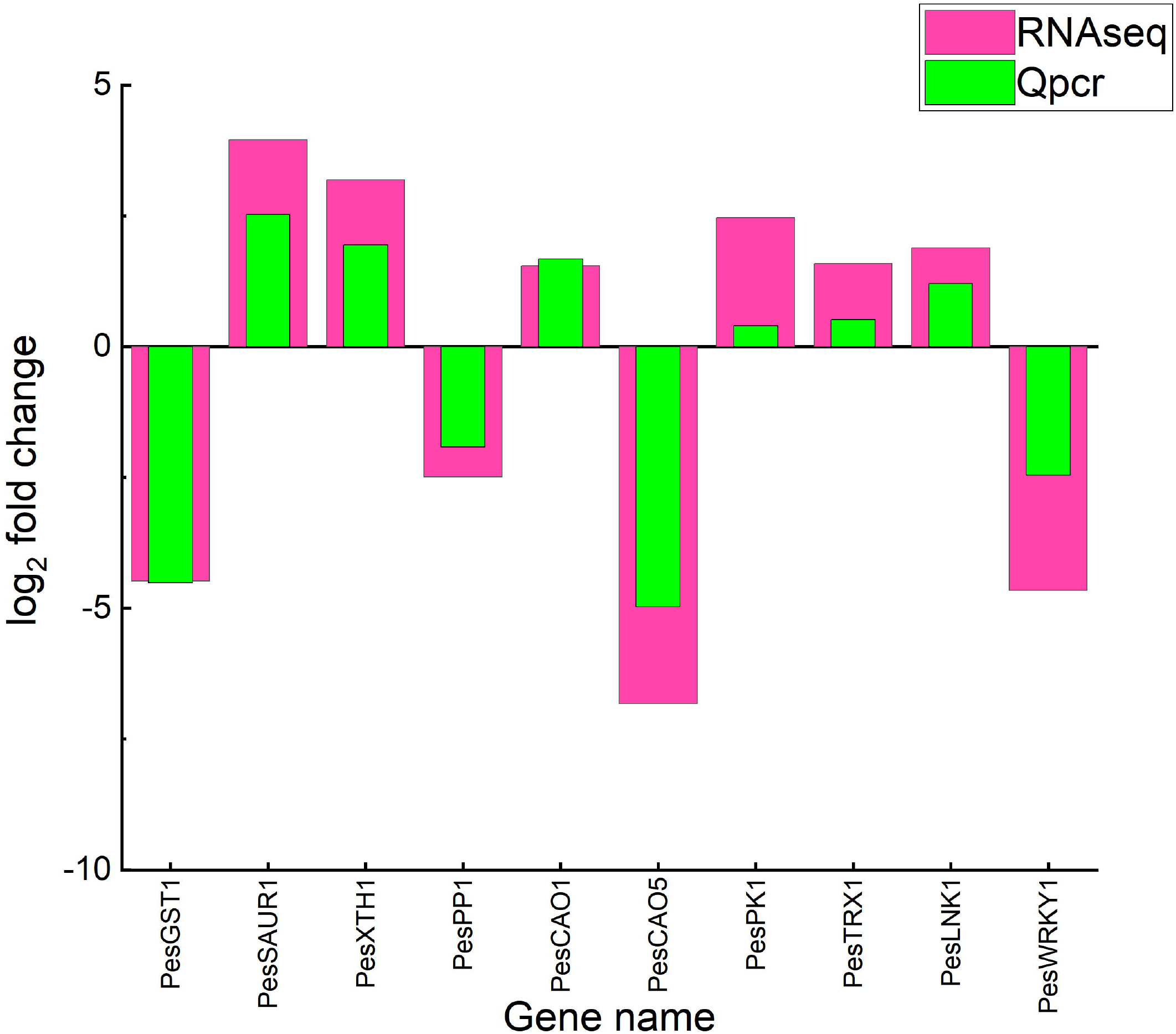

